# Development and Analytical Validation of a Smartphone-Based Quantitative Lateral Flow Immunoassay for Serum Cystatin-C

**DOI:** 10.64898/2026.06.21.733583

**Authors:** Ye Lian, Ruiying Zheng, Chenqi Yang, Leyang Luo, Na Zhang, Guojun Lian, Baoqing Li

## Abstract

Cystatin-C is an important renal function biomarker, and conventional quantification requires centralized laboratory analyzers, which limits timely testing in primary care and resource-limited settings. To address this need, we developed and validated a simple, rapid, and quantitative smartphone-based (SP) lateral flow immunoassay (LFIA) for measuring serum Cystatin-C. The SP-LFIA platform consists of a colorimetric LFIA strip and a custom SP reader with uniform LED illumination and macro lens for image capture. Quantitative image analysis of the colorimetric signal is performed by a dedicated application using a pre-defined third order polynomial calibration model. Following systematic optimization, the assay demonstrated a wide quantitative range of 0.32–8.00 mg/L, with a limit of detection of 0.15 mg/L. Analytical validation conducted according to CLSI guidelines showed excellent precision, with intra- and inter-assay coefficients of variation below 10%, and no significant interference from bilirubin, triglycerides, hemoglobin, or rheumatoid factor. Accelerated stability testing confirmed robust strip performance after storage at 50 °C for 28 days. Method comparison using 100 clinical serum samples showed high agreement with a commercial PETIA reference method (R² = 0.993) and minimal bias. These results indicate that the developed smartphone-based LFIA provides a reliable, cost-effective, and practical tool for point-of-care Cystatin-C monitoring.

## 1. Introduction

Chronic kidney disease (CKD) represents a major and growing global public health burden, affecting more than 10% of the adult population worldwide and contributing substantially to morbidity, mortality, and healthcare expenditure [1–3]. Early identification of renal dysfunction is critical because timely intervention can slow disease progression, reduce cardiovascular complications, and improve long-term outcomes in patients, particularly in the elderly and other high-risk populations, e.g. those with diabetes or cardiovascular disease [4–7]. However, chronic kidney disease (CKD) often has few or no symptoms during early stages, emphasizing the need for a sensitive accessible biomarker for prompt detection and routine monitoring of renal function.

Cystatin-C is an endogenous biomarker that can reliably assess renal function, and estimate glomerular filtration rate (GFR). Unlike serum creatinine, Cystatin-C production is relatively independent of muscle mass, age, sex, and dietary intake, making it particularly useful in risk stratifying in the elderly as well as in patients with diabetes or altered muscle metabolism [2,6–9]. Many clinical studies have demonstrated that elevations of Cystatin-C are associated with adverse renal and cardiovascular outcomes even when GFR estimates based on creatinine appear to be normal [6–8,10–12]. In addition, Cystatin-C has predictive value in determining disease progression and mortality in multiple patient cohorts, including seniors and patients with multiple chronic medical problems [13,14].

Despite the clinical usefulness of Cystatin-C measurement and the increased availability of methods to measure Cystatin-C, routine Cystatin-C measurement remains greatly limited in many health care settings. Currently, the clinical assays available to measure Cystatin-C utilize particle-enhanced turbidimetric or nephelometric immunoassays (PETIA/PENIA) requiring centralized laboratory analyzers, trained personnel, and strict quality control procedures [15–18]. Consequently, the availability of Cystatin-C assays is generally limited in primary care clinics, emergency departments, community hospitals, and resource-limited environments. In addition, the costs associated with laboratory-based testing, as well as the longer turnaround times to obtain results, severely limit the potential for routine monitoring and screening of large populations for Cystatin-C.

Alternative analytical methods for the detection of Cystatin-C have been explored to address these shortcomings, including enzyme-linked immunosorbent assays (ELISA), electrochemical sensors, and lateral flow immunoassays (LFIA) [19–23]. LFIAs offer unique advantages over other methods (i.e., ELISA and electrochemical sensors) in their simplicity, fast analysis, and low cost, making them attractive for point-of-care testing (POCT) applications. However, these traditional LFIAs are primarily qualitative or semi-quantitative in nature and typically require visual interpretation of results or a simple card reader for reading. Such methods introduce operator-dependent variability into the results and make them unsuitable for accurate clinical quantification [22–25].

Recent advances in smartphone-based diagnostic technologies have created new opportunities to improve the quantitative capability of colorimetric LFIAs. Smartphones are capable of providing high-resolution images and have standardized data processing and flexible computer processing capability providing the means to quantitatively analyze signals without specialized instruments [26–30]. Furthermore, when colorimetric LFIA assays are optimally designed and combined with robust image processing algorithms, smartphone assisted LFIAs exhibit improved analytical performance and reproducibility, while still maintaining the operational simplicity and low-cost benefits associated with traditional LFIA methods.

This study was conducted to develop and validate a smartphone-based quantitative colorimetric LFIA for measuring Cystatin-C in serum. By integrating a conventional gold nanoparticle-based LFIA strip with a customized smartphone reader and dedicated image analysis software, the proposed platform seeks to overcome the limitations of traditional LFIA systems while retaining their inherent advantages. The analytical performance, clinical applicability, and agreement with an established PETIA reference method were systematically evaluated using human serum samples. This work positions the developed platform as a practical and accessible solution for decentralized renal function assessment and point-of-care Cystatin-C testing.

## 2. Materials and Methods

### 2.1. Materials

Chicken egg yolk immunoglobulin Y (IgY) and polyclonal goat anti-chicken IgY were purchased from Solarbio Biotechnology (Hangzhou, China). Recombinant human Cystatin-C protein, detection antibody (Cystatin-C Ab1, CY-C2401), and capture antibody (Cystatin-C Ab2, CY-C2402) were obtained from Chuangye Biotechnology (Shaoxing, China). Casein was purchased from Sigma-Aldrich (Shanghai, China). Chloroauric acid (HAuCl₄), boric acid, sodium borate, trisodium citrate, sodium carbonate, sodium azide, and buffer components/stabilizers (Tween-20, sucrose, trehalose, BSA, PBS), were sourced from Sinopharm Chemical Reagent Co. (Shanghai, China). Nitrocellulose membranes (NC140) were supplied by Sartorius (Göttingen, Germany). We obtained the glass fiber conjugate pads (SB08), sample pads (SM3-125), and absorbent pads (ABP-370) from Jinbiao Biotechnology (Shanghai, China).

### 2.2. Instruments

An HM3035 dispensing instrument (Jinbiao Biotechnology, Shanghai, China) was employed for reagent deposition onto the LFIA substrates. The assembled cards were subsequently cut into 3 mm wide strips using a ZQ3500 guillotine cutter (Jinbiao Biotechnology, Shanghai, China). A Redmi Android smartphone (Xiaomi, Beijing, China) equipped with a macro lens attachment (Apexel APL-MS009, focal length 5.5–7 mm) and integrated LED illumination was used as the portable reader for imaging the test strips. The smartphone-macro lens system ensured clear visualization and high-resolution capture of the test line while maintaining a stable and consistent lighting environment for quantitative analysis.

### 2.3. Antibody Conjugation onto AuNPs

Monodisperse gold nanoparticles (AuNPs) were synthesized using the citrate reduction method. Briefly, 100 mL of 0.01% HAuCl4 solution was heated to boiling, followed by rapid addition of 3 mL of 1% trisodium citrate solution under vigorous stirring. The mixture was boiled for an additional 15 min until a wine-red color appeared, indicating AuNP formation. The resultant AuNPs exhibited a characteristic localized surface plasmon resonance (LSPR) peak at 525 nm and had an average diameter of approximately 32 nm, as measured by UV-Vis spectroscopy and transmission electron microscopy (TEM), consistent with previous reports [31].

Antibody conjugation to AuNPs was performed by adjusting two 10 mL aliquots of AuNP (OD 1.0 at 525 nm) colloidal solutions to a final pH of 8.75, using 1% (w/v) K_2_CO_3_; 150 µg Cystatin-C Ab1 was added to one aliquot, and 120 µg IgY was added to the second aliquot, with gentle mixing and incubating for 20 minutes at room temperature. Unoccupied binding sites were blocked by adding 4 mL of 1% BSA in 25 mM borate buffer (pH 8.75) to each mixture and incubating for an additional 20 min. The suspensions were then centrifuged (8000× g, 30 min, 4℃) to collect Cystatin-C Ab1-AuNP and IgY-AuNP pellets, which were subsequently washed three times with 0.25% BSA in 25 mM borate buffer (pH 8.75). Prior to storage at 2–8℃, the conjugates were resuspended in a storage buffer composed of 50 mM sodium borate (pH 8.75), 1% BSA, 0.2% Tween-20, 2% sucrose, 2% trehalose, and 0.2% NaN_3_.

### 2.4. Buffer Solutions

The buffer solutions used in this study were as follows: antibody-conjugate blocking buffer (25 mM borate buffer, 1% BSA, pH 8.75), antibody-conjugate washing buffer (25 mM borate buffer, 0.25% BSA, pH 8.75), conjugate storage buffer (50 mM sodium borate, 1% BSA, 0.2% Tween-20, 2% sucrose, 2% trehalose, 0.2% NaN_3_, pH 8.75), test/control line dispensing buffer (10 mM PBS, 1% sucrose, pH 8.75), sample pad buffer (0.1 M PBS, 1% BSA, 0.05% Tween-20, pH 7.4), conjugate pad buffer (0.1 M PBS, 0.05% Tween-20, 20% sucrose, 1% BSA, pH 7.4), washing buffer (PBS, 0.05% Tween-20), and assay running buffer (PBS, 0.1% Tween-20, 1% BSA, pH 7.4).

### 2.5. LFIA Strip Assembly

LFIA strip assembly involved the integration of four main components: a glass fiber sample pad (SM3-125), a glass fiber conjugate pad (SB08) preloaded with AuNP conjugates, a nitrocellulose membrane (NC140), and an absorbent pad (ABP-370). The test line consisted of Cystatin-C Ab2 (1.2 mg/mL), while the control line contained goat anti-chicken IgY (1.0 mg/mL), both diluted in 10 mM PBS with 1% sucrose (pH 8.75). Test and control lines were striped onto the nitrocellulose membrane at a density of approximately 1 μL/cm using an HM3035 dispenser. Subsequent drying was carried out at 37℃ for 2 h.

The sample pad was pretreated with sample pad buffer (0.1 M PBS, 1% BSA, 0.05% Tween-20, pH 7.4) for 5 min and dried overnight at room temperature. The conjugate pad was immersed in conjugate pad buffer (0.1 M PBS, 0.05% Tween-20, 20% sucrose, 1% BSA, pH 7.4) and dried overnight; after drying, it was dipped in the optimized Cystatin-C Ab1-AuNP and IgY-AuNP conjugate mixture and dried at 37℃ for 1 h. Finally, all components were assembled on an adhesive backing card with 2 mm overlapping between adjacent pads and cut into 3 mm wide strips using a programmable guillotine cutter. The strips were stored at 4℃ in sealed aluminum foil pouches with desiccant until use (Figure 1).

**Figure 1.**
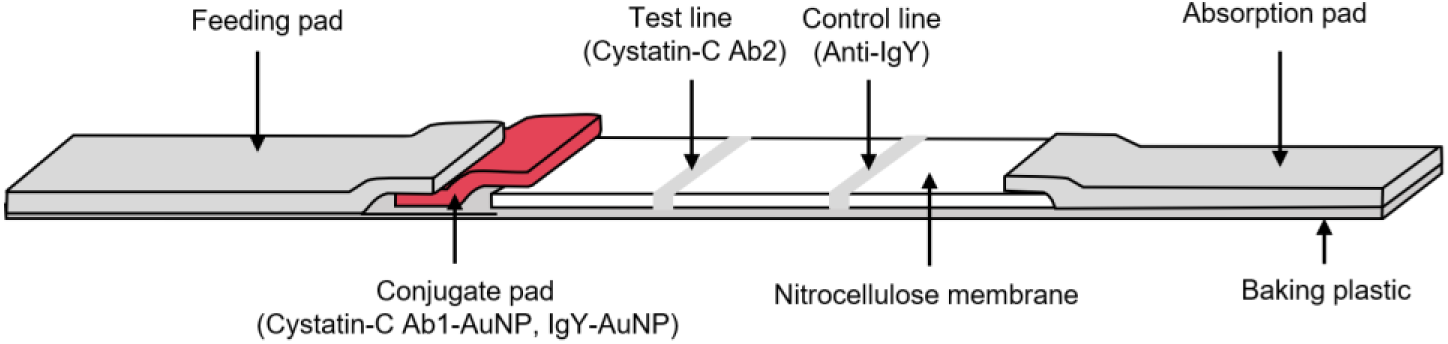
Diagram of LFIA strip components and their spatial arrangement.

### 2.6. Software Development

A custom Android application (“Smartphone rapid detection software for Cystatin C”, version 1.0) was developed to control the smartphone-based reader and perform image analysis. This software is registered with the China National Copyright Administration (Registration No. 2025SR1844772). The application was developed following basic Android app development principles, utilizing Intents for Activity transitions and data transfer to enable user-App interactions. Fragment and ViewPager components were employed for page reusability.

The application consists of two main modules. The first module is the image acquisition module, which uses the smartphone camera equipped with a macro lens and LED ring light to capture images of LFIA strips. A custom SurfaceView implementation provides real-time camera preview for photography function, while an OverlayView class extending View uses Canvas to draw a transparent rectangular window on the preview, creating a constrained photography frame that guides users to align the test line within the designated imaging region. Dynamic permissions for camera and file read/write operations are requested at runtime following Android best practices, integrating with SurfaceView to achieve photography preview function.

The second module is the quantification module, which employs the OpenCV library (version 4.8.0, downloaded from https://opencv.org/releases/) for image processing. Captured images are first resized to 50 × 150 pixels using **I**mgproc.resize(), and then converted to grayscale using Imgproc.cvtColor() with COLOR_BGR2GRAY. The test line, control line, and background regions of interest (ROIs) are identified according to the fixed strip geometry within the reader. The average grayscale values for the test line (I_T_), control line (I_C_), and background (I_B_) are then calculated, and the T/C ratio is computed as:

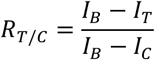

A predefined third-order polynomial calibration model (*y* = *ax*^3^ + *bx*^2^ + *cx* + *d*) embedded in the application is subsequently used to convert T/C ratios into Cystatin-C concentrations. The smartphone application provides users with simple and intuitive methods to capture LFIA strip images and automatically obtain the corresponding Cysta-tin-C values. Images with obvious positioning errors, severe blur, or incomplete strip insertion were excluded from quantitative analysis.

### 2.7. Assay Procedure

To initiate each test, a mixture of 30 µL serum and 80 µL running buffer (PBS, 0.1% Tween-20, 1% BSA, pH 7.4) was prepared and deposited onto the sample pad of the LFIA strip. After the introduction of the sample, the LFIA strip was inserted into the smartphone reader so that the test line was aligned with the imaging area of the smartphone camera. Driven by capillary flow, Cystatin-C in the sample bound to the Cystatin-C Ab1-AuNP conjugates on the conjugate pad, forming Cystatin-C Ab1-AuNP-Cystatin-C complexes. These immunocomplexes were specifically captured at the test line by immobilized anti-Cystatin-C antibody (Ab2), yielding a visible red band with intensity proportional to Cystatin-C concentration. Unbound Cystatin-C Ab1-AuNP and IgY-AuNP continued to migrate, with IgY-AuNP captured by goat anti-chicken IgY at the control line, confirming proper flow. After 15 min of development, the strip was imaged under consistent illumination using the smartphone reader. The custom application analyzed the test line intensity and calculated the Cystatin-C concentration using the predefined third-order polynomial calibration model described above. A schematic of the assay workflow and image analysis is presented in Figure 2.

**Figure 2.**
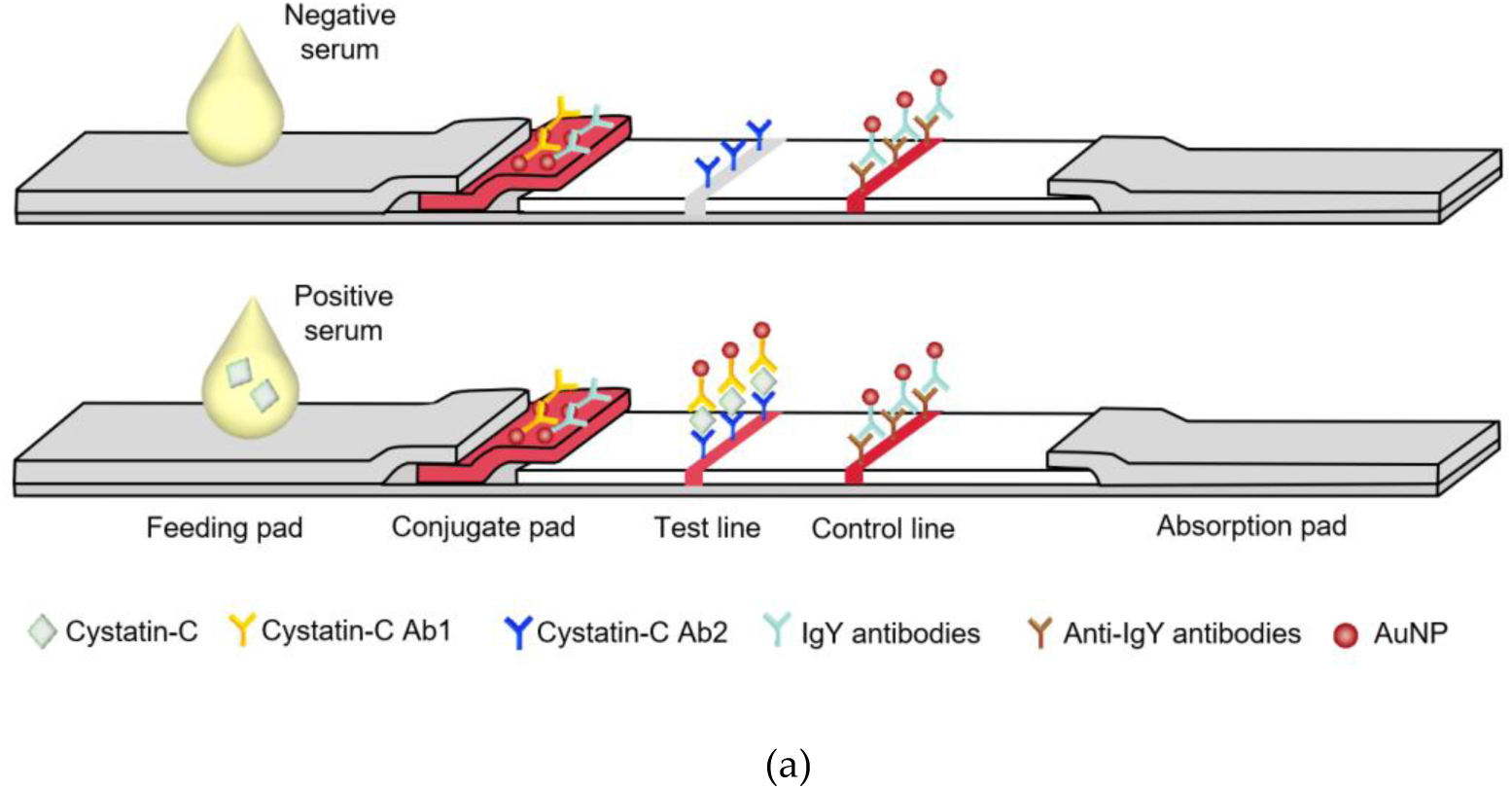
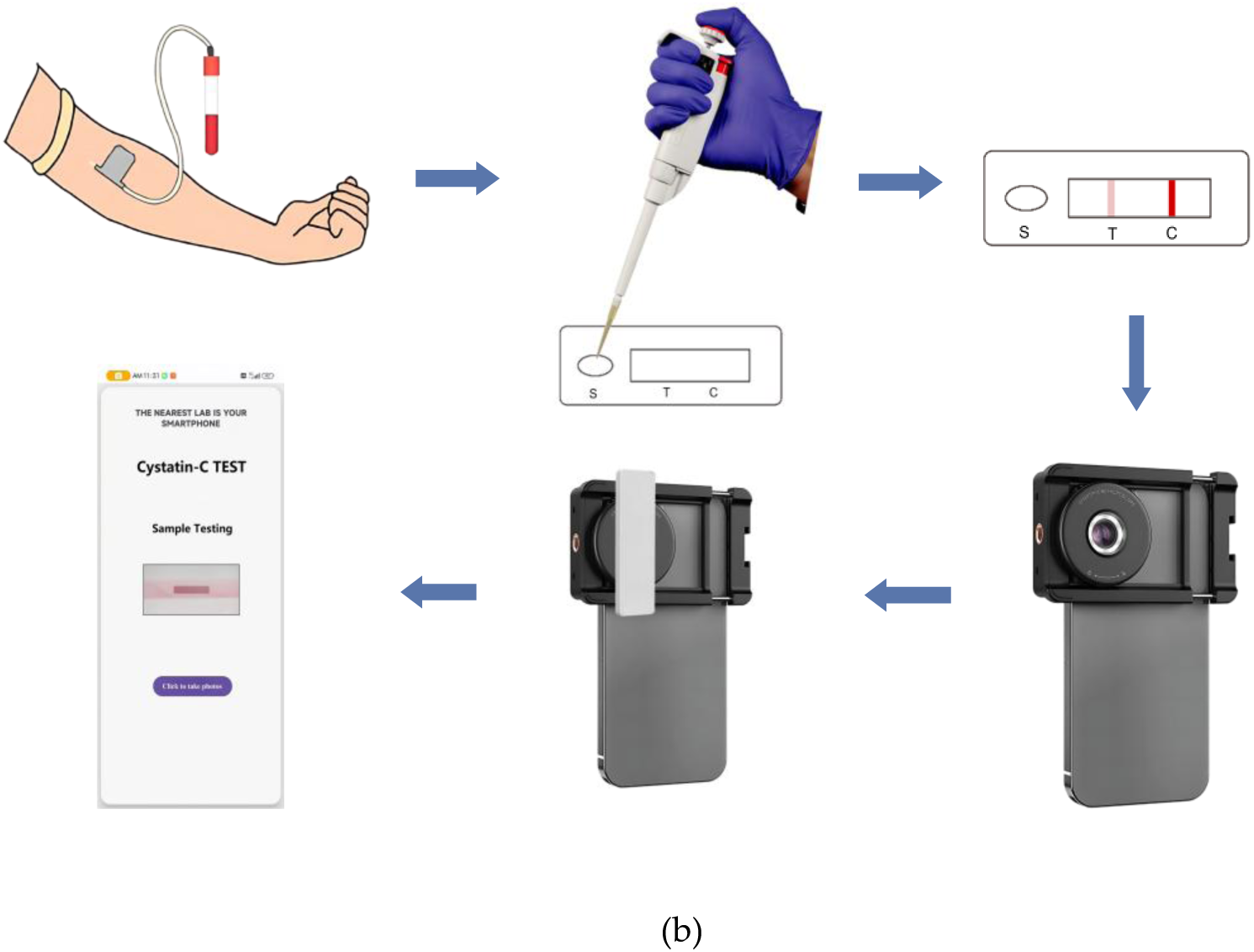
Schematic representation of the testing procedure using the smartphone-based LFIA. (a) The diluted sample migrates through the conjugate pad, where it binds to Cystatin-C Ab1-AuNP conjugates, and continues to the test line (T). There, the complexes are captured, forming a red band whose intensity is proportional to the Cystatin-C concentration. The control line (C) captures IgY-AuNP complexes to confirm valid flow. (b) The strip is inserted into the smartphone reader, and the built-in app analyzes the test line intensity to output the Cystatin-C concentration.

### 2.8. Optimization Procedures

#### 2.8.1. Capture Antibody Concentration Optimization

Capture antibody (Cystatin-C Ab2) was diluted to 0.4, 0.8, 1.2, and 1.6 mg/mL in 10 mM PBS (pH 8.75) containing 1% sucrose. Each solution was dispensed onto nitrocellulose membranes at 1 μL/cm using an HM3035 dispenser. Membranes were dried at 37℃ for 2h. Test strips were assembled and evaluated with Cystatin-C standards at 0.5, 2.5, and 8.0 mg/L (n = 3 per concentration). After 15 min development, strips were imaged using the smartphone-based reader, and T/C ratios were calculated as described in Section 2.5.

#### 2.8.2. Detection Antibody Labeling Amount Optimization

Cystatin-C Ab1 at quantities of 50, 100, 150, and 200 μg was added to 10 mL aliquots of AuNP solution (OD = 1, pH 8.75). Conjugation, blocking, washing, and storage were performed as described in Section 2.3. Conjugate pads were prepared with each labeling amount and assembled into test strips. Evaluation with Cystatin-C standards (0.5, 2.5, and 8.0 mg/L) and T/C ratio calculation were performed as above (n = 3 per concentration).

#### 2.8.3. BSA Concentration in the Running Buffer Optimization

Running buffers containing 0.5%, 1%, 2%, 3%, and 5% BSA in PBS with 0.1% Tween-20 (pH 7.4) were prepared. Test strips (n = 5 per condition) were tested with blank (0 mg/L) and 2.5 mg/L Cystatin-C standards. T/C ratios were determined after 15 min development.

#### 2.8.4. Tween-20 Concentration in the Running Buffer Optimization

Running buffers containing 0.01%, 0.05%, 0.1%, 0.2%, and 0.5% Tween-20 (with 1% BSA in PBS, pH 7.4) were prepared. Test strips (n = 10 per condition) were tested with 2.5 mg/L Cystatin-C standard. T/C ratios and coefficients of variation (CV%) were calculated.

#### 2.8.5. Assay Time Optimization

Test strips were tested with Cystatin-C standards at 0.25, 0.5, 1.0, 2.0, 4.0, and 8.0 mg/L. Images were captured at 5, 8, 10, 12, 15, and 20 min post-sample application (n = 5 per time point). T/C ratios were plotted against time for each concentration.

### 2.9. Validation of the Developed LFIA

After establishing the assay design, a comprehensive analytical validation was conducted, encompassing evaluations of precision, linearity, dynamic range, limit of detection (LOD), limit of quantitation (LOQ), interference testing, comparison with a reference method, and accelerated stability assessment.

#### 2.9.1. Precision

The precision of the Cystatin-C measurement assay was evaluated in accordance with the CLSI EP15-A2 guidelines [32]. Precision was evaluated across a range of Cystatin-C concentrations using serum specimens at 0.75 (low), 3.00 (medium), and 6.00 (high) mg/L. Each specimen was tested 20 times in a single day to estimate within-run precision (repeatability), while each specimen was tested a total of 12 separate days to estimate between-run precision (intermediate precision). For each concentration level, the mean, standard deviation, and coefficient of variation (CV) were calculated.

#### 2.9.2. Linearity and Sensitivity

To determine both the linearity of the assay performance and the sensitivity of the assay, Cystatin-C standard solutions covering the expected clinical concentration range (0 – 8.00 mg/L) were generated and analyzed using the LFIA. The test data were analyzed in accordance with the CLSI EP6-A guidelines [33]. A calibration curve was generated using the predefined third-order polynomial model integrated in the smartphone application, and the quantitative working range of the dose–response relationship was identified. For sensitivity assessment, the LOD was defined as the mean blank signal plus three standard deviations, based on 10 replicate blank measurements, while the LOQ was determined as the lowest concentration yielding a coefficient of variation ≤20%, in accordance with CLSI EP17-A2 guidelines [34].

#### 2.9.3. Interference Testing

Following CLSI EP7-A2 guidelines [35], potential interference from endogenous and exogenous substances was systematically examined. Moderate-level Cystatin-C samples (∼2.50 mg/L) were spiked with common interferents, including bilirubin (0.23 mg/mL), triglycerides (12 mg/mL), hemoglobin (6 mg/mL), and rheumatoid factor (750 IU/mL), and then analyzed using the LFIA. Results from spiked samples were compared against un-spiked controls, with recoveries within ±15% of the control value defined as the absence of significant interference.

#### 2.9.4. Method Comparison

One hundred clinical serum samples were prospectively collected from the Second Affiliated Hospital of Wenzhou Medical University, Wenzhou, China. All samples were obtained with informed consent, and the study protocol was approved by the Ethics Committee of the Second Affiliated Hospital of Wenzhou Medical University (Protocol No. MR-33-26-014646, approval date: February 25, 2026). These specimens that were used for assay validation were obtained from randomly selected patients and were found to represent Cystatin-C concentrations ranging from 0.45 to 7.57 mg/L, providing assay results representative of both low (normal) and high Cystatin-C concentrations. Comparison between the developed LFIA and a commercial particle-enhanced turbidimetric immunoassay (PETIA) reference method for Cystatin-C measurement was then performed in accordance with CLSI EP9-A2 guidelines [36]. The reference method employed a Cystatin C Assay Kit (latex-enhanced immunoturbidimetric assay) from Zhejiang Kuaye Biotechnology Co., Ltd., China, and all measurements were carried out according to the manufacturer’s instructions. The LFIA result for each sample was compared with the corresponding PETIA result to evaluate the agreement between the two methods. Correlation was assessed using linear regression (Pearson’s R²), and agreement was further evaluated with Bland–Altman analysis to determine the mean bias and 95% limits of agreement.

#### 2.9.5. Accelerated Stability

Accelerated stability testing was performed to estimate the potential shelf-life of the LFIA strips. Batches were stored at 21 ℃, 37 ℃, and 50 ℃ for up to 28 days, with samples removed at defined time points (weeks 0, 1, 2, 3, and 4) for evaluation. At each interval, strips were tested using Cystatin-C–positive samples at three concentration levels (1.00, 2.50, and 5.00 mg/L), and the resulting T-line signal intensities were compared with those obtained from fresh strips stored at 2–8 ℃. Stability was defined as acceptable when the signal recovery remained within 85–115% of the initial value. The accelerated study was used to provide a preliminary assessment of strip robustness under stressed storage conditions; real-time stability studies are required to establish the shelf-life under routine ambient storage conditions.

### 2.10. Application of the Developed LFIA to Clinical Serum Samples

The developed smartphone-based LFIA was applied to the determination of Cystatin-C in human serum samples. For each sample, 30 μL of serum was mixed with 80 μL of running buffer and applied to the sample pad of the LFIA strip. After 15 min of development, the strip was inserted into the smartphone reader, and the Cystatin-C concentration was automatically calculated by the custom application using the pre-established calibration model. Each serum sample was also analyzed using the reference particle-enhanced turbidimetric immunoassay (PETIA) for method comparison. All measurements were performed according to the optimized assay conditions described above.

### 2.11. Statistical Analysis

Statistical evaluation of the obtained results was performed using the Statistica package (13.3). Quantitative data are presented as mean ± standard deviation (SD). Linear regression analysis was used to evaluate the relationship between the smartphone-based LFIA and the reference PETIA method. Method agreement was further assessed using Bland–Altman analysis, and the mean bias with 95% limits of agreement was calculated. A paired t-test was used to assess whether the mean difference between the two methods was statistically significant. A p value < 0.05 was considered statistically significant.

## 3. Results

### 3.1. Optimization of Assay Parameters

To establish the assay conditions that yielded optimal analytical performance, the principal parameters affecting the smartphone-based lateral flow immunoassay were systematically optimized using the T/C ratio as the response metric. The T/C ratio was selected because it minimizes the influence of background fluctuation and inter-strip variation, thereby providing a more robust basis for condition screening. Following the optimization strategy commonly adopted in lateral flow assay development, the concentration of capture antibody on the test line, the labeling amount of detection antibody on AuNPs, the BSA content in the running buffer, the Tween-20 concentration in the running buffer, and the assay development time were sequentially evaluated. The optimized conditions are summarized in Figure 3 and Tables 1 and 2.

**Figure 3.**
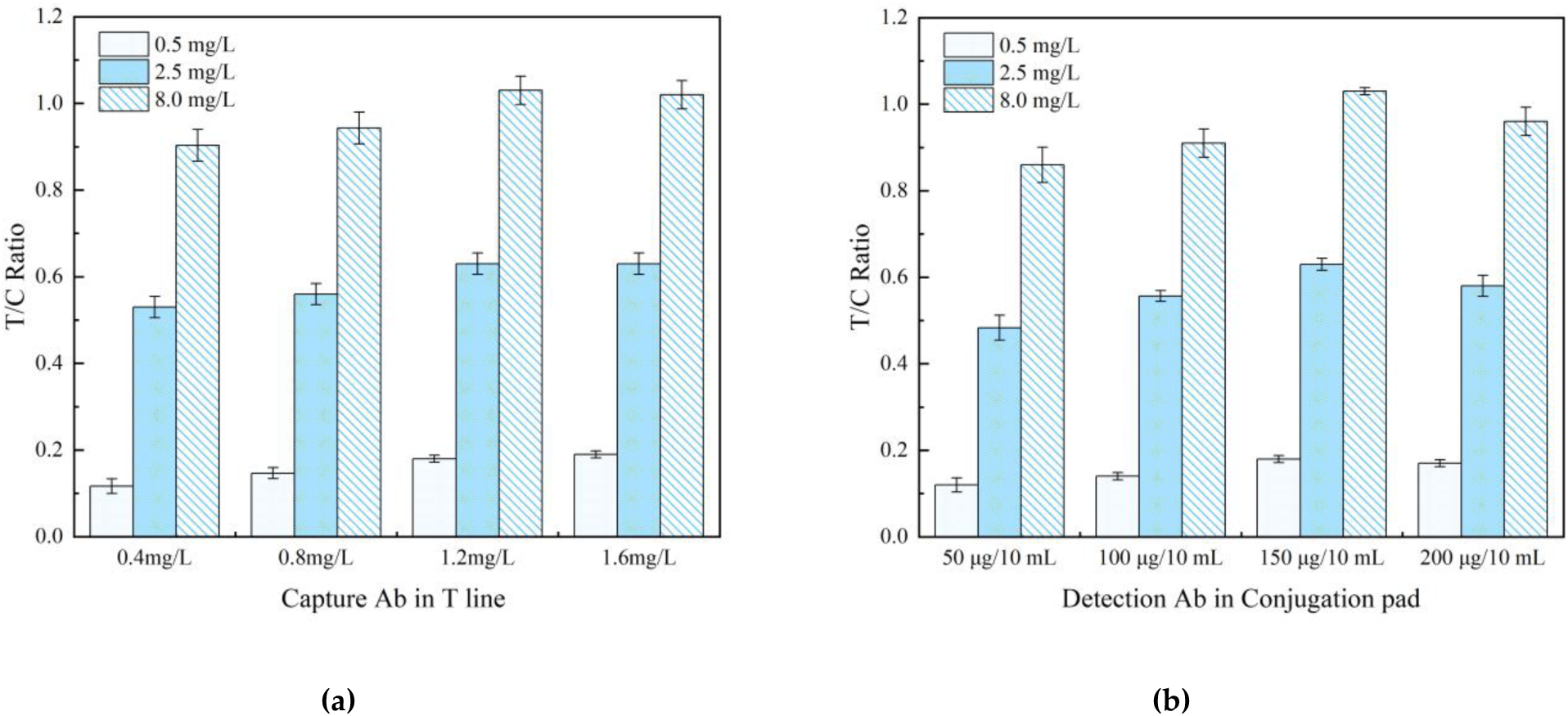
Optimization of capture antibody concentration and detection antibody labeling amount for the smartphone-based LFIA. (a) Optimization of the capture antibody concentration on the test line. The T/C ratio was measured at four Cystatin-C Ab2 concentrations (0.4, 0.8, 1.2, and 1.6 mg/mL) using Cystatin-C standards at 0.5, 2.5, and 8.0 mg/L. (b) Optimization of the detection antibody labeling amount on AuNPs. The T/C ratio was measured using Cystatin-C Ab1 labeling amounts of 50, 100, 150, and 200 μg per 10 mL AuNP solution with the same Cystatin-C standards. In both experiments, the optimal conditions were selected based on signal intensity and assay performance. Data are presented as mean ± SD (n = 3).

**Table 1.** Effect of BSA concentration on T/C ratio.

| BSA Concentration (%) | Blank Sample (0 mg/L) | 2.5 mg/L Standard |
| --- | --- | --- |
| 0.5 | 0.05 ± 0.02 | 0.48 ± 0.04 |
| 1.0 | 0.02 ± 0.01 | 0.62 ± 0.03 |
| 2.0 | 0.02 ± 0.01 | 0.55 ± 0.03 |
| 3.0 | 0.01 ± 0.01 | 0.47 ± 0.04 |
| 5.0 | 0.01 ± 0.01 | 0.37 ± 0.05 |

**Table 2.** Effect of Tween-20 concentration on T/C ratio and reproducibility.

| Tween-20 Concentration (%) | T/C Ratio (0.5 mg/L) | CV% (n = 10) |
| --- | --- | --- |
| 0.01 | 0.12 ± 0.02 | 15.4 |
| 0.05 | 0.15 ± 0.02 | 9.7 |
| 0.10 | 0.18 ± 0.01 | 5.2 |
| 0.20 | 0.16 ± 0.01 | 8.2 |
| 0.50 | 0.13 ± 0.02 | 12.1 |

#### 3.1.1 Capture Antibody Concentration on the Test Line

The concentration of capture antibody immobilized on the nitrocellulose membrane plays a critical role in determining assay sensitivity. As shown in Figure 3A, the T/C ratio increased progressively as the concentration of Cystatin-C Ab2 increased from 0.4 to 1.2 mg/mL for all tested standards. When the concentration was further increased to 1.6 mg/mL, no further significant enhancement in signal was observed. Accordingly, 1.2 mg/mL was selected as the optimal capture antibody concentration, as it provided strong signal intensity without unnecessary reagent consumption.

#### 3.1.2 Detection Antibody Labeling Amount

The amount of detection antibody conjugated to AuNPs was subsequently optimized because it directly affects probe activity and signal generation. As shown in Figure 3B, the T/C ratio increased markedly as the amount of Cystatin-C Ab1 increased from 50 to 150 μg per 10 mL AuNP solution. The highest responses were obtained at 150 μg, whereas a decline in signal was observed when the labeling amount was further increased to 200 μg. For the low-concentration standard (0.5 mg/L), the 150 μg condition also produced the strongest response, indicating improved sensitivity in the low-concentration range. Therefore, 150 μg per 10 mL AuNP solution was selected as the optimal labeling amount for subsequent experiments.

#### 3.1.3 BSA Concentration in the Running Buffer

The concentration of BSA in the running buffer was optimized to achieve an appropriate balance between suppression of nonspecific binding and maintenance of specific signal generation. As summarized in Table 1, the highest T/C ratio for the 2.5 mg/L Cystatin-C standard was obtained at 1% BSA (0.62 ± 0.03), while the blank sample remained low (0.02 ± 0.01). At 0.5% BSA, the blank T/C ratio increased to 0.05 ± 0.02, suggesting insufficient blocking. In contrast, further increasing the BSA concentration to 2–5% resulted in a progressive decline in T/C ratio for the positive sample. Based on these results, 1% BSA was selected as the optimal concentration in the running buffer.

#### 3.1.4 Tween-20 Concentration in the Running Buffer

Tween-20 concentration was further optimized because surfactant content strongly affects sample wetting, flow uniformity, and assay reproducibility. As summarized in Table 2, the highest T/C ratio for the 0.5 mg/L standard was obtained at 0.10% Tween-20 (0.18 ± 0.01), which also yielded the lowest coefficient of variation (5.2%). At 0.01% Tween-20, the T/C ratio was lower and the CV was markedly higher, suggesting poor wetting and less uniform migration. At concentrations above 0.10%, the T/C ratio gradually decreased. Therefore, 0.10% Tween-20 was selected as the optimal surfactant concentration for the running buffer.

#### 3.1.5 Optimal Assay Time

The development of the test line signal was monitored over time (5–20 min) across a range of Cystatin-C concentrations. For high concentrations (>2.00 mg/L), the test line became visible within 8 minutes. Signal intensity increased progressively and reached a plateau before 15 minutes for all concentrations tested, with no significant change observed thereafter. Therefore, 15 minutes was established as the optimal assay time, ensuring complete sample migration and signal stabilization while maintaining rapid analysis (Figure 4).

**Figure 4.**
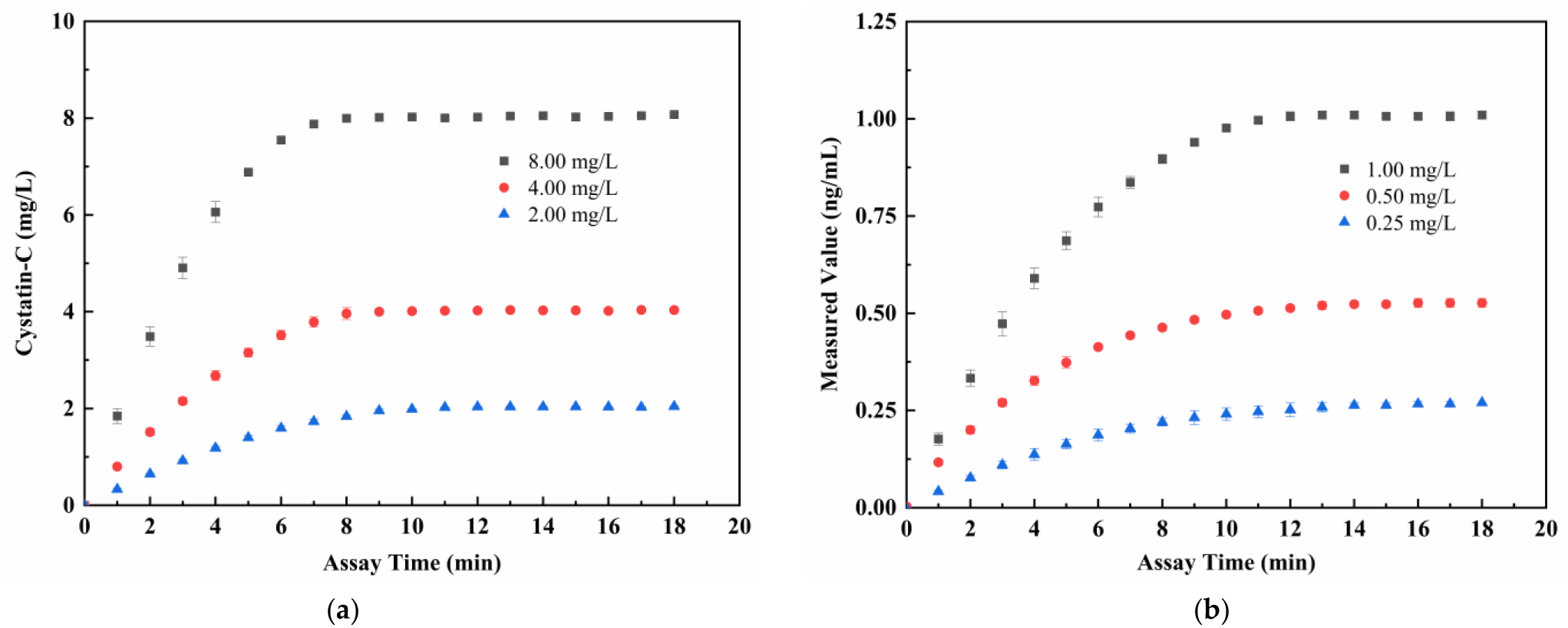
The optimization of the assay time for six Cystatin-C concentrations: (a) 8.00, 4.00, and 2.00 mg/L; (b) 1.00, 0.50, and 0.25 mg/L.

### 3.2. Method Validation

#### 3.2.1 Precision of the LFIA

The precision of the smartphone-based LFIA was evaluated at three clinically relevant Cystatin-C concentrations (low: ∼0.75 mg/L, medium: ∼3.00 mg/L, high: ∼6.00 mg/L). The results, summarized in Table 3, demonstrated excellent repeatability and intermediate precision. The within-run coefficients of variation (CVs, n=20) were 7.40%, 5.12%, and 3.87% for the low, medium, and high levels, respectively. The between-run CVs (n=12) were 9.56%, 6.79%, and 5.02%, respectively. All CV values were well below the 15% acceptance threshold, confirming the high reproducibility of the assay. The raw data for within-run and between-run precision are provided in Table S1 and S2.

**Table 3.** Reproducibility of the smartphone-based LFIA for Cystatin-C.

| Cystatin-C Levels<br>(mg/L) | Within-Run ($n=20$) | | Between-Run ($n=12$) | |
| --- | --- | --- | --- | --- |
|  | Average (mg/L) | CV (%) | Average (mg/L) | CV (%) |
| High (~6.00) | 6.22 | 3.87 | 6.19 | 5.02 |
| Medium (~3.00) | 3.03 | 5.12 | 2.94 | 6.79 |
| Low (~0.75) | 0.74 | 7.40 | 0.78 | 9.56 |

#### 3.2.2 Linearity, LOD, and LOQ

Cystatin-C standard solutions were used to establish the quantitative capability of the assay. The smartphone application provided the calibration curve using the embedded third order polynomial model, which provided a dynamic range from 0.32 mg/L to 8.00 mg/L. The linear regression analysis of the comparison between measured and expected values yielded a correlation coefficient (R²) for the Cystatin-C standard concentration versus raw data concentration of 0.9994, demonstrating excellent linearity for the smartphone-based LFIA (Figure 5). The LOD was calculated based on the mean of the 10 blank samples ± 3 SDs to be 0.15 mg/L. The limit of quantification (LOQ), defined as the lowest concentration measurable with a CV ≤ 20%, was established at 0.32 mg/L, where the observed CV was approximately 10%.

**Figure 5.**
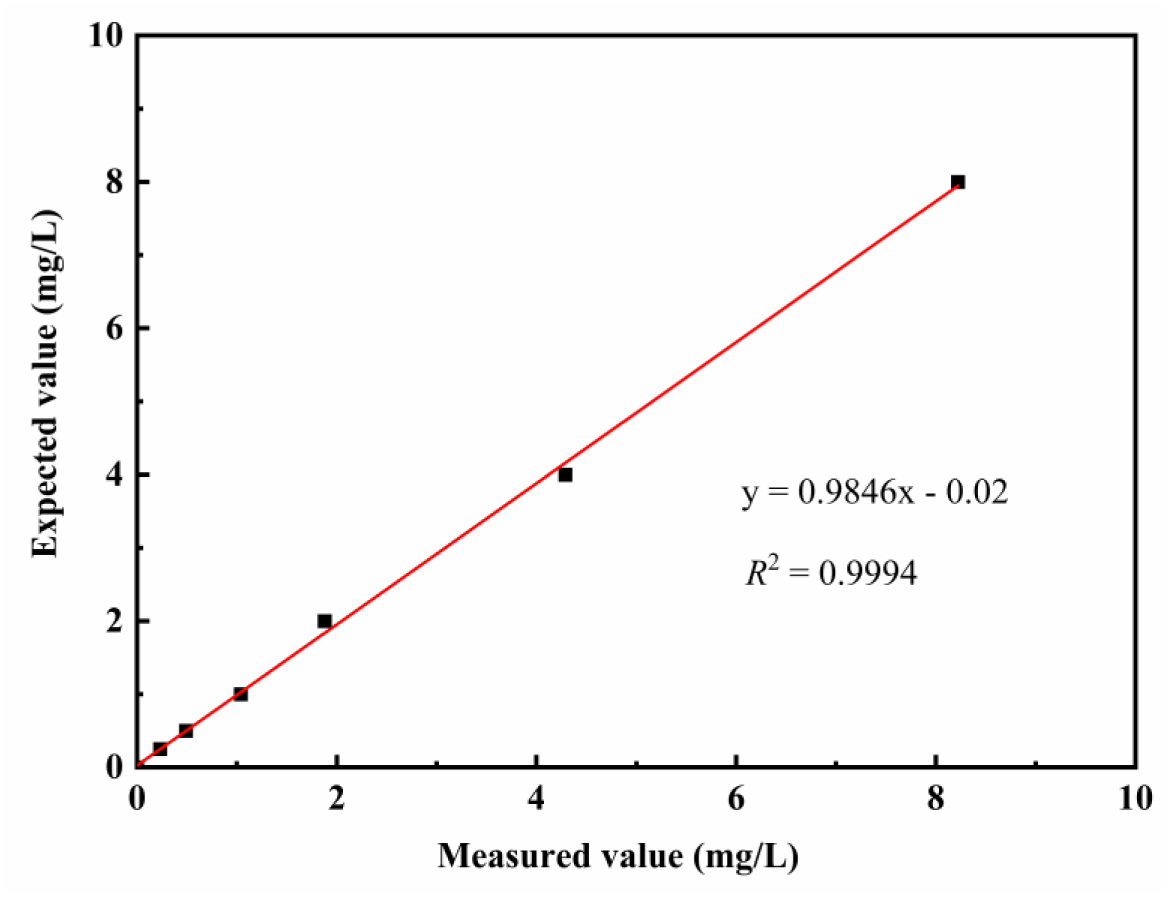
The calibration curve for the developed LFIA was generated using serum calibrators, with each concentration point measured in triplicate.

#### 3.2.3 Interference Testing

The potential interference from common endogenous substances was investigated. As shown in Table 4, the recovery rates for a Cystatin-C sample (2.50 mg/L) spiked with high concentrations of bilirubin, triglycerides, hemoglobin, and rheumatoid factor all fell within the acceptance range of 85–115%. These results demonstrate that the assay is robust and specific, with no significant interference from these substances. Detailed recovery results for each interferent are summarized in Table S3.

**Table 4.** Interference results of the smartphone-based LFIA for Cystatin-C.

| Interferent | Concentration of Interferent | Cystatin-C (mg/L) | LFIA Assay Result (mg/L) | Recovery (%) |
| --- | --- | --- | --- | --- |
| Control | - | 2.50 | 2.57±0.14 | 103 |
| Bilirubin | 0.23 mg/mL | 2.50 | 2.86±0.15 | 115 |
| Triglycerides | 12 mg/mL | 2.50 | 2.87±0.16 | 115 |
| Hemoglobin | 6 mg/mL | 2.50 | 2.87±0.11 | 115 |
| Rheumatoid Factor | 750 IU/mL | 2.50 | 2.12±0.13 | 85 |

#### 3.2.4 Accelerated Stability Testing

The stability of LFIA strips was assessed at 21℃, 37℃, and 50℃ for an extended period of 28 days. The results of the accelerated stability testing show that the test signal values for low (1.00 mg/L), medium (2.50 mg/L), and high (5.00 mg/L) concentration results remained between 85% to 115% of their initial values after being stored for 28 days at each temperature (Figure 6). Therefore, there was no significant deterioration of analytical performance for the LFIA strips during the accelerated storage testing. These findings strongly suggest that LFIA strips have good storage stability. However, to determine product shelf life and confirm long-term stability under normal handling/storing conditions, real-time stability studies are necessary.

**Figure 6.**
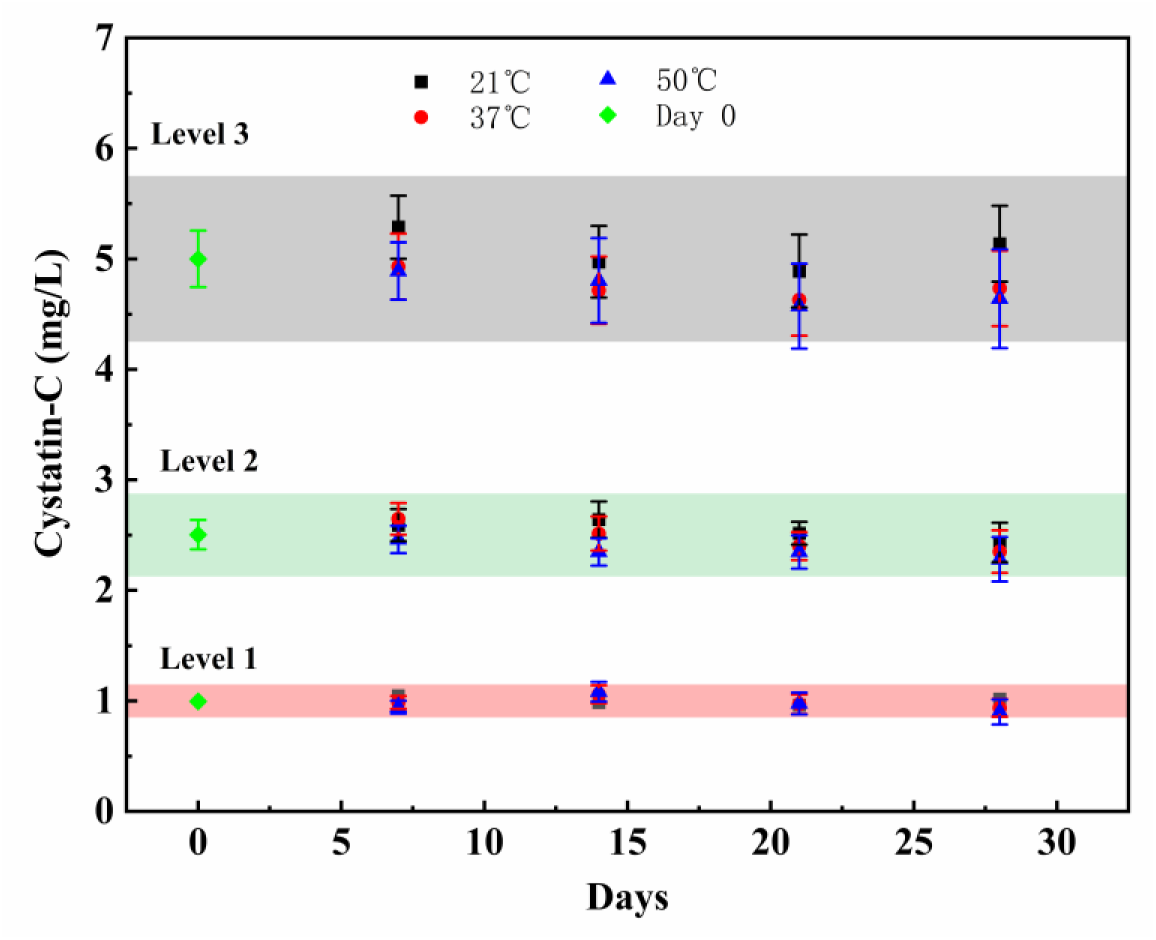
Accelerated stability testing of LFIA strips. Color region indicates data within 85 to 115% across levels.

### 3.3. Application of the Method and Statistical Analysis of the Results

The Cystatin-C measurement performance of the smartphone-based LFIA was evaluated using human serum specimens, and the analytical performance results were compared to the commercially available PETIA assay. To establish the practical utility of the proposed method under clinical conditions, 100 clinical serum samples with a wide range of Cystatin-C concentrations were analyzed.

Each serum sample was tested using the optimized lateral flow immunoassay (LFIA) method. The sample concentration of Cystatin-C was automatically calculated with respect to the calibration curve by a custom smartphone application. All samples were also tested with the manufacturer-authorized PETIA method. After both analytical techniques were complete, paired results were compared statistically through linear regression analysis.

Linear regression analysis indicated a very strong correlation between LFIA and PETIA results. As illustrated in Figure 7, the regression equation was y = 0.996x − 0.014, yielding an R-squared value of 0.993. This indicates very good agreement between the LFIA and PETIA methods within the clinically relevant range of Cystatin-C concentrations.

**Figure 7.**
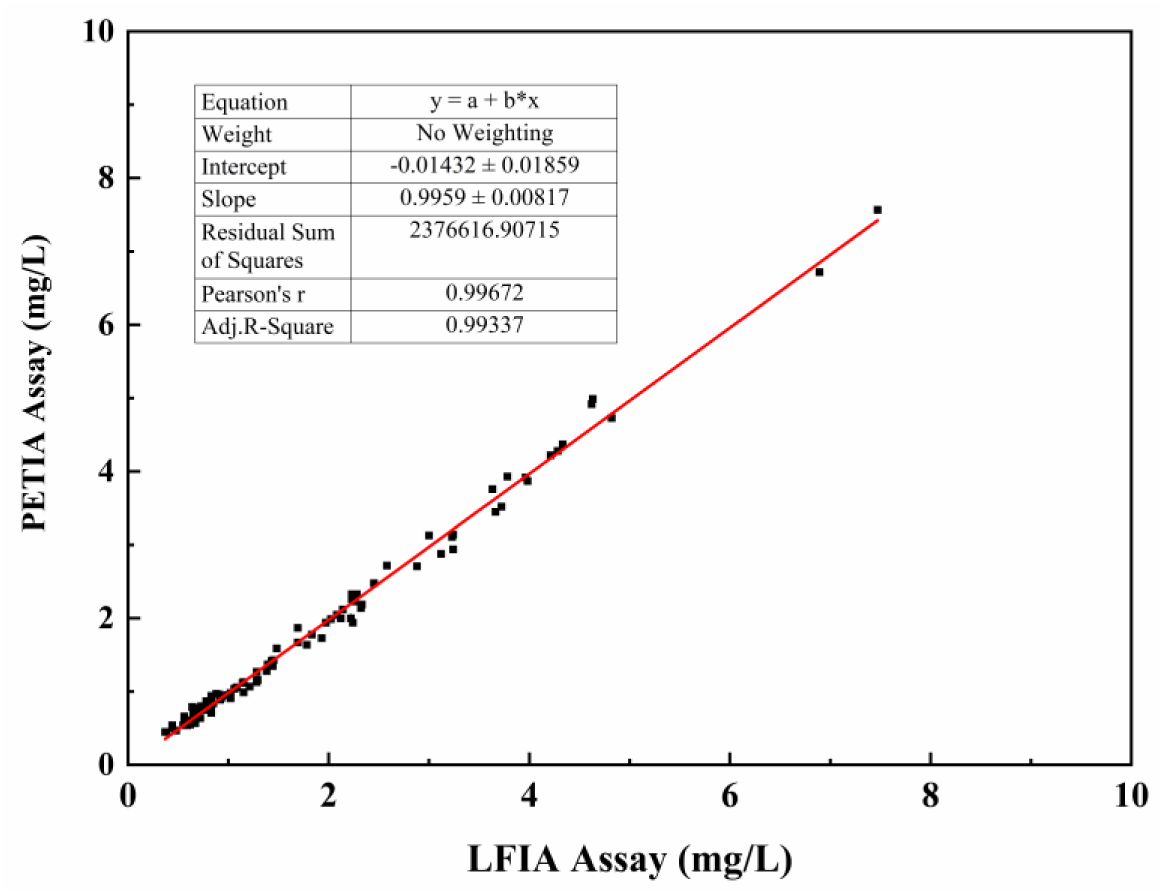
Linear correlation between Cystatin-C concentrations measured by the smartphone-based LFIA and the PETIA reference method using 100 human serum samples. The solid line represents the linear regression fit (*y = 0.996x − 0.014*, *R*² = 0.993), indicating excellent agreement between the two analytical methods across the clinically relevant concentration range.

To further assess method agreement, Bland–Altman analysis was performed (Figure 8). The mean bias between the LFIA and PETIA methods was −0.02 mg/L, indicating minimal systematic difference between the two assays. The 95% limits of agreement ranged from −0.24 to 0.20 mg/L, and the mean difference was not statistically significant (paired t-test, p = 0.0583). No systematic trend was observed across the measured concentration range, supporting good agreement between the two methods within the tested clinical interval.

**Figure 8.**
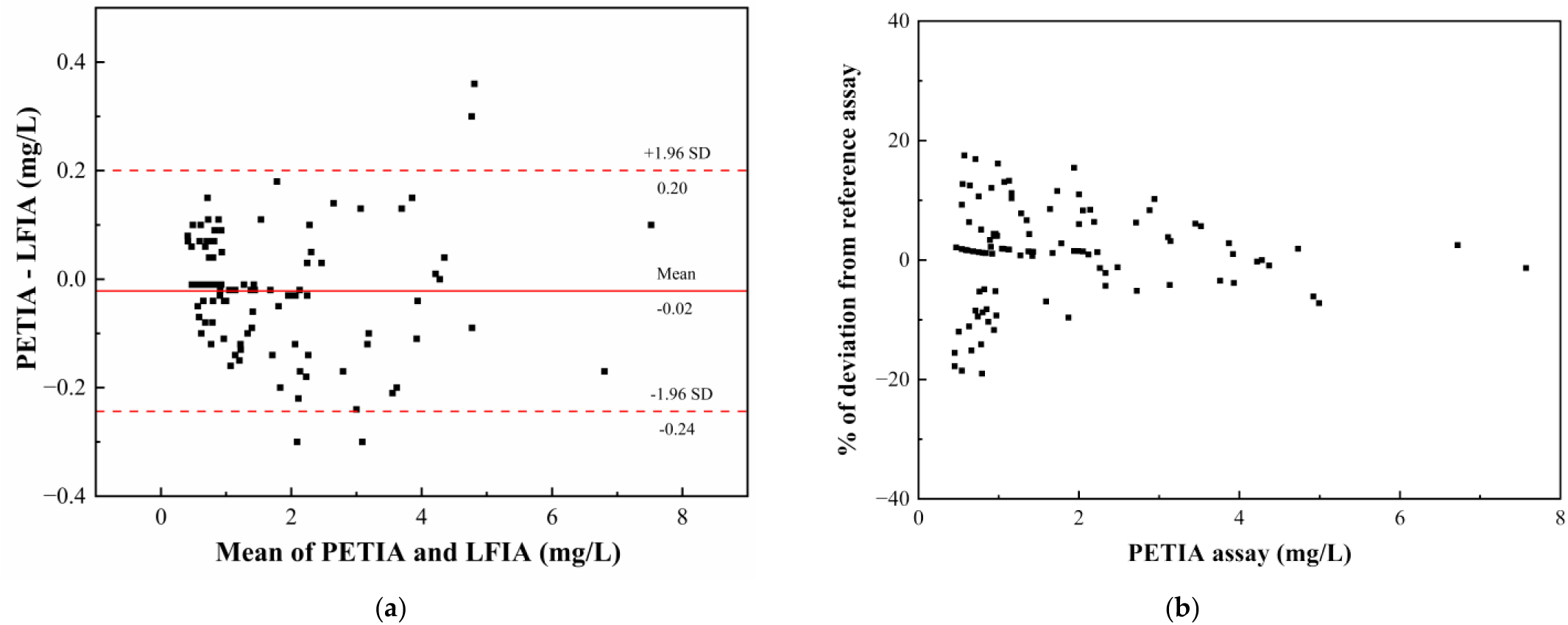
Bland–Altman analysis of LFIA versus PETIA results. **(a)** Differences between methods plotted against their averages, showing the mean bias (solid line) and the 95% limits of agreement (dashed lines). The mean bias was −0.02 mg/L, and the 95% limits of agreement ranged from −0.24 to 0.20 mg/L. **(b)** Percentage differences relative to PETIA values across the measured Cystatin-C concentration range (0.45–7.57 mg/L).

## 4. Discussion

Cystatin-C has been identified as an accurate and sensitive biomarker for assessment of renal function and to identify early indications of renal impairment due to its relative independence to muscle mass, age, and dietary factors relative to creatinine-based markers. Accurate and timely cystatin C measurements are therefore essential for screening and risk stratification in patients with early chronic kidney disease (CKD) [6–9]. However, traditional methods for quantifying Cystatin-C have depended primarily on automated immunoassays conducted in centralized laboratories (particle-enhanced turbidimetric and nephelometric methods) that require trained personnel, automated analyzers, and extensive turnaround times for completion [15–18], which severely limits accessibility to these immunoassays in community healthcare settings, emergency rooms, and resource-poor environments.

In this investigation, we developed a smartphone-based LFIA for quantitative analysis of serum Cystatin-C levels. Results support the conclusion that the proposed method had comparable analytical performance to laboratory-based immunoassays but better characteristics with regards to speed, portability, and simplicity of use. By integrating a conventional colorimetric LFIA strip with a customized smartphone reader and a dedicated image analysis algorithm, the platform effectively bridges the gap between centralized laboratory testing and decentralized point-of-care diagnostics [21–23,29,37].

The analytical sensitivity and dynamic range of the assay analysis meet clinical requirements for quantifying renal function. The limit of detection is 0.15 mg/L, which can reliably identify CKD below the normal reference range in healthy adults [38]. Furthermore, the assay covers a span of 0.32–8.00 mg/L and eliminates or reduces the need for dilution of samples, simplifying routine patient management procedures. These performance characteristics support the suitability of the assay for both screening and followup applications in clinical practice.

In the method comparison involving 100 clinical serum samples utilizing LFIA and PETIA, the smartphone-based LFIA achieved excellent agreement with the commercial PETIA reference method. Linear regression analysis achieved a near-unity slope and high R-squared value. Bland–Altman analysis confirmed there were no systematic differences between the two methods and only minimal mean bias across the measured concentration range [36]. These findings illustrate that the LFIA is comparable or similar to the established laboratory methods, suggesting potential for decentralized analysis of Cystatin-C. However, broader multicenter validation is still warranted before routine clinical implementation.

Comparative performance is summarized in Table 5. The present study suggests that the smartphone-based colorimetric LFIA has an excellent balance of sensitivity, dynamic range, and practical accessibility. The limit of detection (0.15 mg/L) is lower than that of Zhang et al. [0.98 mg/L] and comparable with that of Bikkarolla et al. (0.18 mg/L) [22,38]. A major distinguishing factor of this work is that the image analysis system is integrated into the smartphone, therefore eliminating subjectivity in measurement interpretation compared to previous LFIA studies [22,38,39].

**Table 5.**
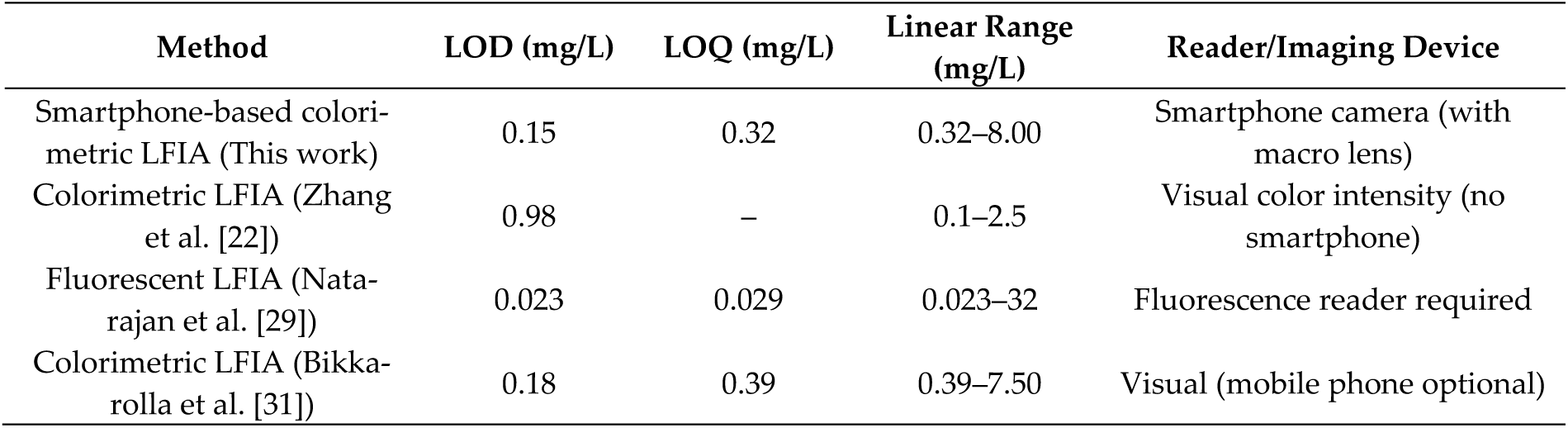
Performance comparison of cystatin-C detection methods between this work and literature.

| Method | LOD (mg/L) | LOQ (mg/L) | Linear Range (mg/L) | Reader/Imaging Device |
| --- | --- | --- | --- | --- |
| Smartphone-based colorimetric LFIA (This work) | 0.15 | 0.32 | 0.32–8.00 | Smartphone camera (with macro lens) |
| Colorimetric LFIA (Zhang et al. [22]) | 0.98 | – | 0.1–2.5 | Visual color intensity (no smartphone) |
| Fluorescent LFIA (Natarajan et al. [29]) | 0.023 | 0.029 | 0.023–32 | Fluorescence reader required |
| Colorimetric LFIA (Bikkarolla et al. [31]) | 0.18 | 0.39 | 0.39–7.50 | Visual (mobile phone optional) |

Advanced platforms such as the fluorescent LFIA described by Natarajan et al. [29] may be superior, with limits of detection in the range of 0.023 mg/L and extended linear range; however, have limited scalability and routine application in decentralized and/or limited-resource situations due to the equipment costs of fluorescence readers [26]. In contrast, the present study used colorimetric gold nanoparticles for labeling in order to maintain simplicity and reduce costs. The limit of detection is also clinically relevant for CKD screening and routine measurements of renal function.

Potential sources of variability in measurement results exist with colorimetric LFIA systems. Variability in the equalization of membrane properties, efficiency of antibody immobilization, stability of conjugated gold nanoparticles, and sample matrix effects (e.g., hemolysis or lipemia) may introduce variability in measurement results [21–23]. Standardized strip production and consistent illumination and imaging through specialized smartphone readers will minimize these sources of variability. Results from the third-order polynomial calibration curve, along with standardized acquisition of images and analysis of grayscale values using defined regions of interest, will minimize variability in measurement associated with uneven color distribution along the test line and minimize interpretation errors by users in measurement results [29,37]. However, improper sample selection or handling will produce variability in measurement results, thus highlighting the need for standardized operating procedures for future clinical use.

From a methodological and engineering standpoint, the proposed smartphone-based LFIA represents an equilibrium between analytical performance and system complexity. As compared to fluorescence or lanthanide-based quantitative LFIA systems that are dependent on specialized optical instrumentation [25–27], the present method retains most of the inherent advantages of gold nanoparticle-based methods, such as low cost, robustness, or ease of manufacture, while producing reliable quantitative output from smartphone-assisted evaluation of images. Such a balance will facilitate greater implementation on a large scale and provide viability for decentralized applications in community healthcare.

The newly developed LFIA clinical application has been validated with its ability to provide quick results, low volume sample requirement, and ease of use. All of these features are especially useful for elderly patients, those who have chronic renal failure, frequently monitor their renal function, and patients seen in outpatient offices or in home health care [10–12]. Because they can now quickly determine Cystatin-C levels, patients will be more compliant to treatment and physicians will be able to make timely clinical decisions without relying solely on centralized laboratory facilities.

Future research should focus on validating the current LFIA with specific groups of patients. In particular, further assessment of feasibility using patients suffering from chronic kidney disease, with diabetic nephropathy or the elderly, is warranted because small changes in Cystatin-C levels have significant prognostic value [6–9,13,14]. Longitudinal studies that look at disease progression and response to therapy would further enhance the clinical utility of the LFIA. Additionally, integration of the LFIA with telemedicine platforms and electronic records will facilitate remote monitoring of Cystatin-C levels, data-driven clinical decision making for patients, and the broader use of decentralized testing for renal function.

This study has multiple limitations even though it has shown promise with the analytical comparison and clinical comparison of the LFIA system. The first limitation is that this study was done using samples from one site and should be confirmed with a larger number of sites before broad clinical implementation of the method. Secondly, this study did not examine the performance of the smartphone system across multiple makes/models of smartphones and new photographic conditions as part of the validation procedure. The third limitation is that the estimated shelf life of the LFIA device was based upon accelerated stability testing of gold nanoparticles and has yet to be confirmed with longterm real-time studies. All three of these considerations need to be elucidated in future studies before wider use of the LFIA technology can occur in a clinical setting.

## 5. Conclusions

In this study, a smartphone-based quantitative colorimetric lateral flow immunoassay was successfully developed and validated for the determination of Cystatin-C in human serum. By integrating a conventional gold nanoparticle-based LFIA strip with a customized smartphone reader and dedicated image analysis software, the proposed platform enables reliable quantitative analysis while maintaining the inherent advantages of simplicity, low cost, and rapid operation.

The results of this study for the Cystatin-C assay show that the LFIA developed had a low limit of detection, broad range of quantitative values, and good correlation with a commercial PETIA method for the Cystatin-C serum specimens. These results for the smartphone-enhanced LFIA provide accurate and reproducible Cystatin-C measurements with the operational advantages of ease of use, low-cost testing, and rapid analysis. Overall, this work highlights the potential of smartphone-assisted colorimetric LFIA as a practical platform for decentralized and point-of-care Cystatin-C testing. Further multicenter clinical validation and real-time stability studies are warranted before routine clinical implementation.

## Supplementary Materials

The following supporting information can be downloaded at: [https://zenodo.org/records/19216545]. Table S1: Raw data for the coefficient of variation for the within-run assay; Table S2: Raw data for the coefficient of variation for the between-run assay; Table S3: Raw data for interference testing.

## Author Contributions

Conceptualization, Y.L. and R.Z.; Methodology, Y.L., R.Z. and C.Y.; Software, Y.L., C.Y., and L.L.; Validation, Y.L., R.Z. and N.Z.; Formal Analysis, N.Z. and L.L.; Investigation, Y.L., R.Z. and C.Y.; Resources, G.L. and B.L.; Data Curation, Y.L. and R.Z.; Writing—Original Draft Preparation, Y.L. and R.Z.; Writing—Review & Editing, G.L., B.L. and N.Z.; Supervision, G.L. and B.L.; Funding Acquisition, G.L. and B.L. All authors have read and agreed to the published version of the manuscript.

## Institutional Review Board Statement

The study was conducted in accordance with the Declaration of Helsinki, and approved by the Ethics Committee of The Second Affiliated Hospital of Wenzhou Medical University (Protocol No. 2026-K-51-01, approval date: January 30, 2026).

## Informed Consent Statement

Informed consent was obtained from all subjects involved in the study.

## Data Availability Statement

The data presented in this study are available on request from the corresponding author.

## Acknowledgments

The authors thank all volunteers and clinical staff involved in sample collection and assay evaluation.

## Abbreviations

The following abbreviations are used in this manuscript:

AuNP: Gold Nanoparticle
BSA: Bovine Serum Albumin
CLSI: Clinical and Laboratory Standards Institute
CKD: Chronic Kidney Disease
CV: Coefficient of Variation
CysC: Cystatin C
ELISA: Enzyme-Linked Immunosorbent Assay
GFR: Glomerular Filtration Rate
IgY: Immunoglobulin Y
LFIA: Lateral Flow Immunoassay
LOD: Limit of Detection
LOQ: Limit of Quantification
LSPR: Localized Surface Plasmon Resonance
PBS: Phosphate Buffered Saline
PENIA: Particle-Enhanced Nephelometric Immunoassay
PETIA: Particle-Enhanced Turbidimetric Immunoassay
POCT: Point-of-Care Testing

